# *In vitro* FLASH irradiation of A549 lung cancer cells and IMR90 healthy human fibroblasts in the synchrocyclotron room of a clinical proton therapy system

**DOI:** 10.1101/2024.11.13.623375

**Authors:** Adrián Zazpe, Inés del Monte-García, Nerea Palao, Fernando Cerrón, Mateo Cueto-Remacha, Paula Linzoain-Agos, Minerva Iniesta-González, Martina Quartieri, Ángel M. Cuesta, Samuel España, Guillermo Velasco, Juan Antonio Vera-Sánchez, Luis Mario Fraile, Alejandro Mazal, José Manuel Udías, Almudena Porras, Paloma Bragado, Álvaro Gutierrez-Uzquiza, Daniel Sánchez-Parcerisa

## Abstract

**Purpose:** This study aims to investigate the FLASH irradiation effect on lung tumor (A549) and healthy fibroblast (IMR90) cell lines using an irradiation station installed at the synchrocyclotron room of a clinical proton facility without any permanent beamline modifications.

**Methods and Materials:** An irradiation system composed of a lead scatterer and 3D-printed positioning system was designed and fabricated to operate within the beamline gap of the IBA Proteus One proton therapy facility after the extraction of the beam from the synchrocyclotron. A dosimetric analysis of the produced irradiation field was carried out using radiochromic films. FLASH and conventional-rate irradiations were conducted on relevant cell lines for lung cancer at the isocenter of the treatment room. Biological assessments post-irradiation included clonogenic and viability assays for cell survival, immunofluorescence analysis of p21 protein expression, and flow cytometry analysis for cell cycle arrest evaluation.

**Results:** The irradiation system successfully delivered homogeneous and repeatable FLASH dose rates (>900 Gy/s) with a positioning accuracy of 1 mm and dose uniformity within 10%. Clonogenic assays revealed no statistically significant differences in survival between FLASH and conventional dose rates for both A549 and IMR90 cell lines, although a trend towards higher viability was observed in IMR90 cells under FLASH conditions. Flow cytometry demonstrated significant differences in cell cycle arrest patterns at doses above 7 Gy, with FLASH-irradiated cells exhibiting a decrease in G2/M phase arrest compared to conventional rates. Immunofluorescence analysis of p21 expression showed no significant differences between irradiation modalities.

**Conclusions:** The developed irradiation station effectively facilitates FLASH radiotherapy experiments in a clinical proton facility, achieving the necessary dose rates without hardware modification or extra tuning of the facility. Our analysis reported notable alterations in cell cycle dynamics suggesting distinct biological responses between FLASH and conventional rates in both healthy and tumor cells. These findings contribute to the emerging understanding of the FLASH effect and support the potential for its differential impact on cancerous versus healthy tissues.

## 1. Introduction

The re-discovery of the dose-rate effect, coining the term FLASH, was published by the Institut Curie in Paris (Favaudon 2014), highlighting a drastic reduction in the probability of causing lung fibrosis in healthy tissue of mice irradiated with electrons at high dose rate (>40 Gy/s), vs. the same dose at conventional rate, while maintaining therapeutic efficacy in tumors. Subsequent studies in animal models have explored responses such as pulmonary fibrosis (Fouillade 2020), neurocognitive involvement (Montay-Gruel 2018) or epithelial necrosis (Vozenin 2019) among others; with electrons (Favaudon 2014), photons (Montay-Gruel 2018) and protons (Diffenderfer 2020).

However, the mechanisms behind the FLASH effect are not yet fully understood (Ahmad 2024). It was initially associated with instantaneous oxygen depletion (Durante 2018), but more recent research suggests that it could be related to a differential ability between healthy and tumor cells to detoxify and repair cellular injury caused by reactive oxygen species (Spitz 2019; Favaudon 2022), to dose-rate dependent changes in lipid peroxidation (Froidevaux 2023, Portier 2024), or to an increased protection of circulating immune cells (Jin 2020). Reported experiments indicate not only a dose rate threshold, but that also a minimum integral dose is needed to observe the FLASH effect.

While the body of data from *in vivo* experiments supporting the FLASH effect is mostly consistent, there is very limited data from *in vitro* experiments (compiled in recent reviews by Adrian 2022 and Friedl 2022), and it does not allow for relevant conclusions to be drawn about the nature of the FLASH effect or the conditions necessary for it to occur.

The reported results vary greatly between endpoints, cell lines, radiation qualities and oxygenation and temperature conditions. For example, a significant decrease (FLASH vs. conventional) in the number of chromosomic aberrations in human blood lymphocytes irradiated with moderate photon doses (2-3 Gy) was reported by two studies carried out over 30 years apart (Prempree 1969, Koriakina 2005), although this finding could not be replicated by a comparable study with electrons (Purrott 1977). A similar decrease in chromosomic aberrations was found in CHO (Chinese hamster epithelial ovary) cells irradiated with FLASH-rate 20-MeV protons vs. conventional rate X-Rays (Schmid 2011), but the dose-rate and LET effects in this case could not be easily decoupled.

When expression of the doble strand break (DSB) associated protein 53BP1 is considered, a differential FLASH effect was reported for IMR90 and MCF5 lung fibroblasts (Fouillade 2020). Nevertheless, this effect could not be observed by the same authors when irradiating A549 (lung cancer) cells, or by other groups irradiating healthy skin fibroblasts AG01522B (Hanton 2019) or cancer cells of lines HNSCC4 (Head and neck squamous cell carcinoma), MDA-MB-231 (breast) or HeLa (cervical) (Adrian 2021). Complexity of the cell damage produced by radiation can also be assessed by cell cycle arrest, and these authors (Adrian 2021) also found no significant differences between FLASH and conventional rates for any of the three cell lines under study. A similar study did find some difference between pulsed (∼10^9^ Gy/s) and continuously irradiated (∼30 Gy/s) protons, with the latter inducing a higher amount of HeLa cells in arresting in the G2/M phase, 10 hours post-irradiation (Auer 2011).

Clonogenic assays have also produced mixed results. A comprehensive study (Adrian 2021) on 7 different cell lines (6 tumoral, 1 healthy) irradiated with FLASH and conventional rate electrons observed a tendency of FLASH rates having a slightly lower killing capacity, but it was found statistically significant for only 4 of the tumoral cell lines, and only for doses above 6 Gy. Similarly, Montay-Gruel (2019) reported on a FLASH-related decrease in the survival of mouse glioblastoma cells irradiated with 20 Gy, but no effect was seen with 10 Gy. Study conditions are indeed relevant: Adrian (2022) recently reported on an observed FLASH effect for melanoma MM576 cells at 9 Gy (electrons), but with remarkable differences for samples seeded pre and post irradiation. These differences were, in fact, higher in magnitude than the reported dose-rate effect. Even one recent study observed an apparent anti-FLASH effect (Ventakesulu 2019), where high-dose rate electrons result in an increased killing capacity for two pancreatic cancer cell lines. Older studies also showed contradictory results: most report isoeffective killing of ultra-high dose and conventional-dose rate irradiations (Zackrisson 1991 in Chinese hamster fibroblasts, Cygler 1994 in glioma and melanoma cells), while one study seems to observe a clear FLASH effect in CHO cells (Michaels 1978), but it does not report on it explicitly.

Finally, oxygenation conditions have proven extremely relevant in modulating the FLASH effect. Three recent studies have observed a FLASH effect only for hypoxic/physioxic conditions (<4% O_2_ pressure): with electrons for DU145 prostate cancer cells (Adrian 2019), with helium ions for A549 and H1437 lung cancer cells (Tessonier 2021) and for electrons in spheroidal 3D models of A549, HT-29 (colorectal adenocarcinoma) and MDA-MB-231 cancer cells (Khan 2021).

Currently, the ability to perform research with proton beams at ultra-high dose rates is limited by the lack of proton facilities able to reach FLASH dose rates. In clinical cyclotrons or synchrocyclotrons, reducing the energy to the desired range causes significant intensity losses, so current (experimental) implementations of FLASH with clinical proton therapy systems use the highest energy of the system combined with either target-specific degraders or shoot-through treatment planning strategies.

At the IBA proteusOne system (Henrotin 2016), there is a gap in the beamline where the beam leaves the vacuum at the exit of the synchrocyclotron and travels in air for about 4 cm, before reentering the vacuum after the energy degrader. In this space, the beam is at its highest energy and intensity and therefore able to produce FLASH dose rates. But there are several challenges associated with using this system: (1) the field size is small and inhomogeneous; (2) the position and optics of the beam are not easily reproducible, as it is situated upstream of the focusing magnets, and (3) it is located inside the accelerator bunker, where activation is high and access to the room is therefore limited.

In this article we present our experience with a portable irradiation system designed for this gap, addressing the aforementioned challenges through a lead scatterer (degrader) designed to increase field size and homogeneity, together with a 3D-printed sample positioning system that facilitates an accurate installation in a short time, minimizing the time spent by experimenters inside the (highly activated) bunker, thereby reducing their radiation exposure.

To test the system in action, we performed a FLASH irradiation of two different cell line samples: lung adenocarcinoma cells (A549) and healthy lung fibroblasts (IMR90). Similar samples were irradiated at conventional dose rate, at the patient treatment room of the facility. Post-irradiation analysis included survival assays, study of the expression of p21 protein via immunofluorescence and study of cell cycle arrest with flow cytometry.

## 2. Materials and methods

### Design and fabrication of the holder

The positioning system was designed to fit exactly in the beamline gap (Figure 1A) and fitted with ruler-type marks allowing for a 1-mm precise positioning. The structure was 3D-printed in PLA and contained a space for a lead scatterer and holders for two Eppendorf vials, where biological samples are placed in two regions of interest (ROIs) of 2 × 5 mm^2^ each (Figure 1B). After an initial study where the optics of the nude beam at the point of interest was determined with radiochromic films, a lead scatterer (shown in Figure 2A) was designed using Monte Carlo simulation in TOPAS (Perl 2012). It was manufactured at our workshop as a 2×3 cm^2^ lead block with maximum thickness of 1.1 cm and a concave inner face. The mean energy with which 230-MeV protons arrive at the samples after traversing the scatterer was calculated to be 204.7 MeV, with a standard deviation of 0.8 MeV.

**Figure 1.**
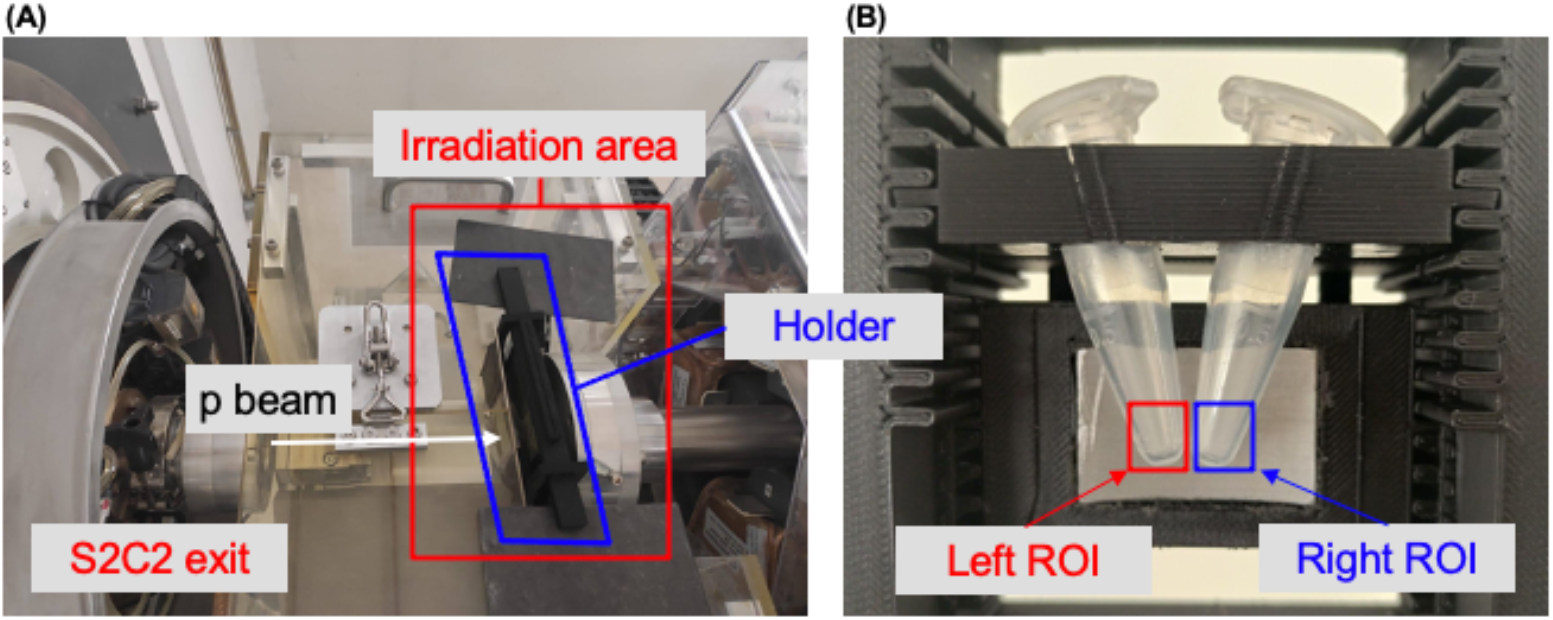
Experimental setup for proton irradiation. (A) View of the irradiation area outside the synchrocyclotron (S2C2) with sample holder in place. (B) View of the 3D-printed holder and scatterer and Eppendorf tubes in place for irradiation.

**Figure 2.**
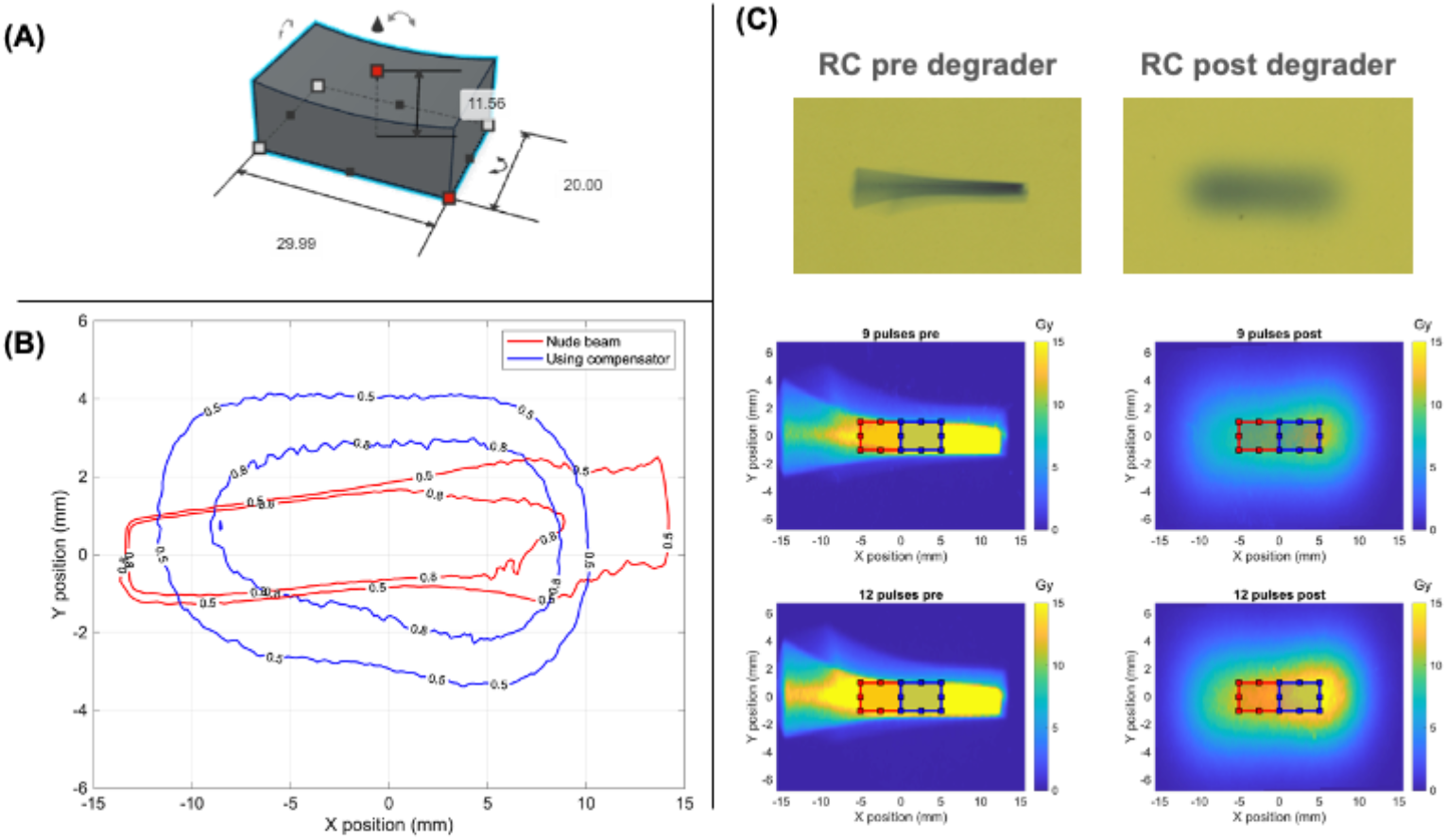
(A) 3D model of the scatterer; (B) 50% and 80% iso dose curves before (red) and after (blue) lead scatterer; maximum local dose values of 2.25 Gy/pulse and 1.30 Gy/pulse before/after scatterer. (C) Scanned radiochromic films and obtained dosimetry for irradiations with 9 and 12 shots, before and after lead scatterers.

### Irradiation

The irradiation system was tested in successive irradiations at the IBA Proteus One facility in physical mode using 230-MeV protons and doses in the 0-12 Gy range. For normal operation, the synchrocyclotron produces 10-μs pulses with a frequency of 1kHz (Mazal 2021, García-Díez 2023). With the collaboration of the IBA personnel, for FLASH irradiations, the system was set to single-pulse mode in physical mode while keeping the beam current at 100 nA, and dose was then controlled by giving a finite number of pulses.

The first irradiation was used to evaluate the dosimetric characteristics of the irradiation field, as well as its repeatability, via radiochromic film dosimetry. Pulse trains with 3, 6, 9 and 12 consecutive pulses (with a separation of 1 ms) were delivered through the scattering block and into the film.

In the second part of the experiment, biological samples were irradiated. One million cells of each line (as described below in “Cell lines and cultures”) were suspended in 1 ml of complete medium in Eppendorf vials, kept in ice, and centrifuged at 1200 rpm for 5 minutes and pelleted right before FLASH irradiation. For these experiments, replicates were also irradiated at conventional rate using a 10×10cm^2^ field at the treatment room and a spread-out Bragg peak with a range of 3 cm and a modulation of 1 cm, using blocks of solid water to ensure a correct coverage of the samples, avoiding dose gradients. Differences in proton energy and modulation for both conditions are further explained in the dosimetry section and further analyzed in the discussion section.

Despite the high activation of the bunker room, the quick setup of the holder allowed dose to experimenters (measured with active-reading online dosimeters and validated with thermoluminescent dosimetry) to remain below 6 μSv for the complete experiment, according to the specific radioprotection protocol designed for this experience. Also, passive gamma and neutron dosimetry for the personnel of the facility did not show any impact with respect to historical records.

### Dosimetry

Previously calibrated EBT-XD films were used for the dosimetry of the irradiated fields, after an on-the-spot recalibration following known protocols (Ruiz-Morales 2020). Films were scanned in transmission using a flatbed scanner Epson 1000XL at a resolution of 150 dpi.

Metrics of beam shape, homogeneity, symmetry and repeatability were evaluated on the dose maps extracted from the films using MATLAB scripts which used a solid image registration algorithm to account for scanning errors in positioning.

To account for different proton energies received by samples at FLASH and conventional rates, RBE(Gy) doses were calculated from physical doses using a phenomenological model (McNamara 2015) using values of 8 Gy and 3 Gy, respectively, for the alpha/beta ratios of cancer and healthy cell lines, with dose-averaged LET values estimated with TOPAS at 0.8 keV/μm and 4.1 keV/μm for entrance-dose and SOBP areas respectively. Further information on the RBE modeling is provided as supplementary data (Figure S1, Table S1).

### Cell lines and cultures

Human adenocarcinoma cell line A549 (CCL-185) was kindly donated by Dr Alcaraz (UB university, Barcelona) and cultured under a humidified atmosphere of 5% CO_2_ at 37ºC in RPMI Media 1640 (Gibco, Waltham, MA), supplemented with 10% fetal bovine serum (FBS, Gibco), 1% penicillin – streptomycin mixture (Gibco) and 0.5% amphotericin B (fungizone) solution (Cytiva). Human healthy lung fibroblast cell line IMR90 (CCL-186) was obtained from ATCC and cultured under a humidified atmosphere of 5% CO_2_ at 37ºC in DMEM Media (Gibco, Waltham, MA), supplemented with 10% fetal bovine serum (FBS, Gibco) and 1% penicillin – streptomycin mixture (Gibco).

### Survival assays

To study the survival of the cell lines after irradiation, two kinds of assays were conducted. Cancer A549 cells were studied using clonogenic assays, whereas for IMR90 cells, lacking clonogenic ability, viability assays were used. The protocol was as follows: right after irradiation, cells were resuspended in normal growth media, seeded and incubated in 6 multi-well plates undisturbed for ten days. A549 cells were seeded in increasing densities with the dose going from 300 to 4,000 cells. The respective cell numbers for reseeding were calculated based on the plating efficiency and expected survival, both calculated from previous in-house experiments. For the IMR90 line, 300 cells were seeded for all conditions. After 10 days, cells were fixed and stained with crystal violet solution in ethanol, For A549 cells, the number of colonies was counted with an automatic script after scanning the plate; for IMR90 cells, the number of remaining viable cells was determined manually using the line-transect method.

### Cell cycle analysis

Right after irradiation, for both cell lines, 330000 cells/condition were seeded in p60 plates and incubated for 48h undisturbed. After that period, they were washed twice with PBS on ice and trypsinized with 0.05% Trypsin-EDTA (Gibco) solution. The suspension was centrifuged at 1200 rpm for 5 minutes at 4ºC and the cells were washed with PBS and fixed with cold ethanol (70%). Resuspended cells in PBS were incubated for 15 minutes with propidium iodide/RNase Staining Buffer (BD Pharmingen #550825) at RT (room temperature). Finally, they were resuspended in PBS and the cell cycle was analyzed by flow cytometry. The raw data obtained from the flow cytometry facility was analyzed with FlowJo. Histograms of the variable FL2–A were analyzed with a MATLAB script to obtain the area under the curve for curves corresponding to the different phases of the cell cycle, as shown in Figure 5A. Cells containing a *2n* DNA content are classified as G0/G1 phases (first peak in the histogram); whereas cells containing a *4n* DNA content are classified as G2/M phases (second peak in the histogram). Cells falling between those two peaks are labeled as phase S. Finally, cells containing a lower amount of DNA than *n* are labeled as SubG0 phase and correspond with apoptotic cells.

### Immunofluorescence analysis of p21

Right after irradiation, for both cell lines, 330000 cells/condition were seeded in 25mm-diameter round glass coverslips and incubated for 48h undisturbed. After that period, they were washed twice with PBS, fixed with 4% para-formaldehyde (PFA) on ice for 20 minutes, washed with PBS, permeabilized with PBS-0.5% Triton X-100 for 5 minutes at RT and PBS-0.1% SDS for another 5 minutes. Then, they were incubated with blocking solution (PBS-3% BSA-1.5% normal goat serum) for 30 minutes at RT.

Next, cells were incubated with anti-p21 (Cell Signaling #2947) antibody (1:100) in blocking solution overnight at 4ºC. After that, the coverslips were washed three times with PBS and incubated with secondary antibody goat anti-rabbit-Alexa 555, 1:500 (Invitrogen A32732) and DAPI (PanReac #A4099, 1:1000) in blocking solution. After washing twice with PBS and once with distilled water, coverslips were mounted with ProLong™ Gold antifade reagent (Invitrogen). Images were captured using a Nikon Eclipse TE300 microscope coupled to a camera. For p21 quantification, the percentage of positive nuclei/field was determined using a self-developed Matlab script.

### Statistical analysis

For the fitting of experimental data to models such as the linear-quadratic (LQ) model (section 3.2) or to equation [1] (section 3.4), with varying uncertainties in dose (X axis) and biological outcome (Y axis), the technique of bootstrapping was used. For each datapoint, 1000 new sets of datapoints were randomly generated from two-dimensional Gaussian distributions around each value and fitted to the models of interest. Error bands in the plots represent the obtained 1-sigma confidence interval for the models. Two-tailed P-values were calculated from the *z* statistic and are shown with stars in the plots: (*) = p<0.05; (**) = p<0.01.

## 3. Results

### 3.1 Dosimetric characterization of the irradiation field

The irradiation system produced a usable irradiation field (after the lead scatterer) of approximately 60 mm^2^ with a positioning accuracy of 1 mm. With doses in the order of 1 Gy per pulse, a pulse duration of about 10 μs (García-Díez 2023) and a pulse repetition time of 1 ms, resulting average dose rates in the sample are in the order of 1000 Gy/s, well over the established FLASH threshold, with instantaneous doses above 10^5^ Gy/s. The measured doses were coherent with the number of protons delivered per shot, as registered in the irradiation logs. Since the beam is clearly asymmetric, the dose rate and dose per pulse in the right ROI is about 18% higher than in the left ROI: calculated values are 0.91 Gy/pulse in the left and 1.08 Gy/pulse in the right, yielding dose rates of 910 Gy/s and 1080 Gy/s, respectively.

Radiobiology studies require homogeneous and repeatable radiation fields in the sample. Therefore, two main sources of uncertainty were assessed. Firstly, dose inhomogeneity within each ROI in the canonical case (assuming a perfect positioning), was assessed, obtaining a value better than 7% in all cases analyzed.

For the latter, we studied mean dose uncertainty within a ROI caused by sample positioning. A bootstrapping study was conducted with 1000 simulated ROI placements, sampled from a 2D gaussian distribution with σ_x_ = σ_y_ = 1 mm. Mean dose relative error was better than 10% for all cases under study (Figure 3A). A third source of uncertainty, variation of field shape between irradiations, was neglected after analysis of the irradiated films: once dose maps from different irradiations were coregistered, their relative dose distributions (measured dose / number of shots) were almost identical in all cases (Figure 3B), ensuring a correct repeatability of the irradiation field.

**Figure 3.**
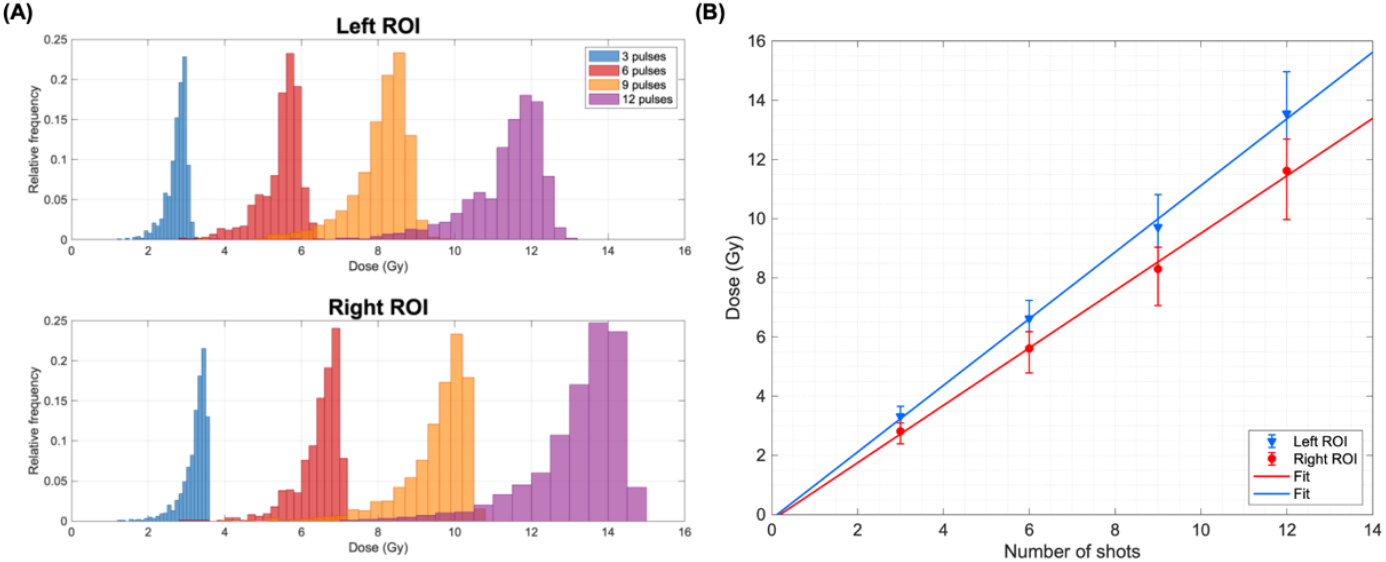
(A) Mean dose histograms obtained for the bootstrapping study (1000 iterations) in left (top) and right (bottom) ROIs. (B) Measured mean dose (error bars show standard deviation) within left and right ROIs as a function of the number of shots delivered.

These combined dosimetric uncertainties were used to determine the error bars in delivered dose for all subsequent biological experiments.

### 3.2 Viability and cell survival

To assess cell survival after irradiation, a clonogenic survival assay was performed for the A549 lung cancer cell line, irradiated in the left ROI of the system. Figure 4A shows the surviving fraction of cells after being irradiated with increasing doses at a conventional (blue) and FLASH dose rates (red), along with their respective fits to the LQ model. As shown in Figure 4A, survival decreased as dose increased in both FLASH and conventional dose rates. Nevertheless, the *Z* test performed on the fitted models showed no significant difference in survival between both dose rates with a 95% confidence level. While the point at D = 12 Gy might indicate a tendency for FLASH rate to be slightly more effective than conventional rate, this difference is also not statistically significant due to the high dosimetric uncertainty in the FLASH arm and steepness of the curves at this point.

**Figure 4.**
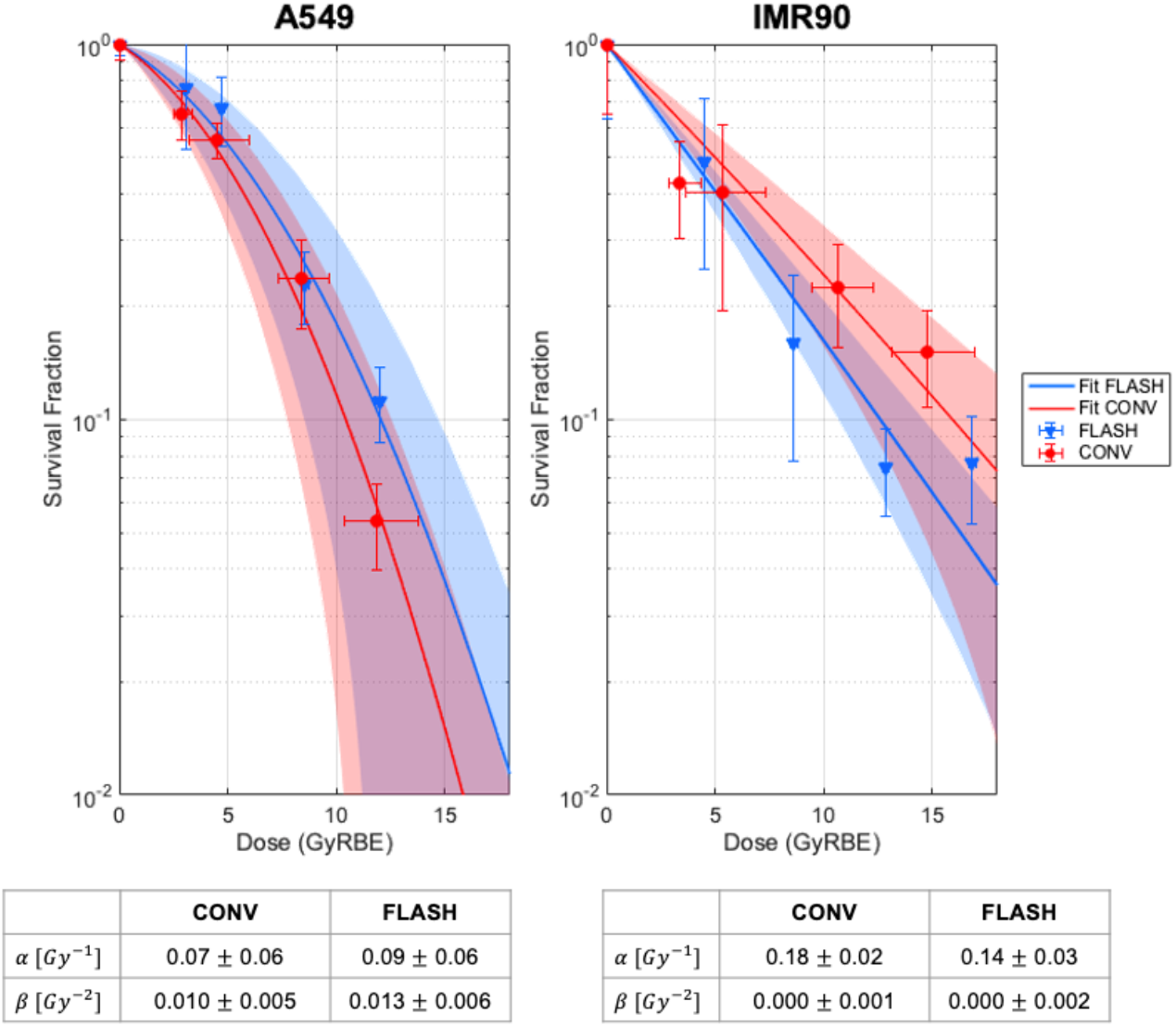
Survival assays at FLASH (red) and conventional (blue) dose rates for lung cancer cell line A549 (A) and healthy fibroblast line IMR90 (B). Shaded areas indicate 1-sigma confidence intervals for the linear-quadratic model fit; tables indicate fit parameters.

For the healthy IMR90 cell line, a viability assay was carried out to assess the long-term effects of irradiation depending on the dose rate. The right ROI of the irradiation device was used. The results of such assay are shown in Figure 4B. A clear tendency for a higher viability in the FLASH branch can be observed, although the *Z* test performed on the fitted models comparing both values of the LQ *α* parameter showed no significant difference between both dose rates at a 95% confidence level (p∼0.27).

### 3.3 Cell cycle arrest

A flow cytometry assay was conducted on both cell lines to evaluate the effect of the dose rate on the cell cycle 48h after irradiation. Figure 5 shows the obtained results from analyzing the histograms of fluorescence intensity: Figure 5A details the histogram analysis process, Figure 5B shows the evolution of the FL2-A histograms with increasing dose for A549 cells at conventional rate. Figures 5C and 5D represent the fraction of cells in each phase of the cycle for A549 and IMR90 cells, respectively. For A549 cancer cells, a significant differential effect was observed at doses higher than 7 Gy, where FLASH induced a lower fraction of cells in arrest in the G2/M phase. These cells were, in turn, in the G0/G1 or S phases. No significant changes in SubG0 phase, were observed for any of the doses at either dose rate, as the fraction of cell fragments was below 1% in all cases under study. For the IMR90 cell line, a similar effect was observed, with significantly fewer cells in the G2/M phase for the FLASH arm for doses above 7 Gy. It is worth noting that since the IMR90 fibroblasts are not proliferative cells, the basal percentage of cells arrested is already above 80% for non-irradiated controls. In this case, the amount of SubG0 (apoptotic cells) is not negligible and increases with the dose, with a significantly higher fraction of SubG0 cells for the FLASH arm for doses above 10 Gy. For both cell lines, the unirradiated controls have slightly different cell cycle distributions between the two arms, but the dose-dependent effects are evident.

**Figure 5.**
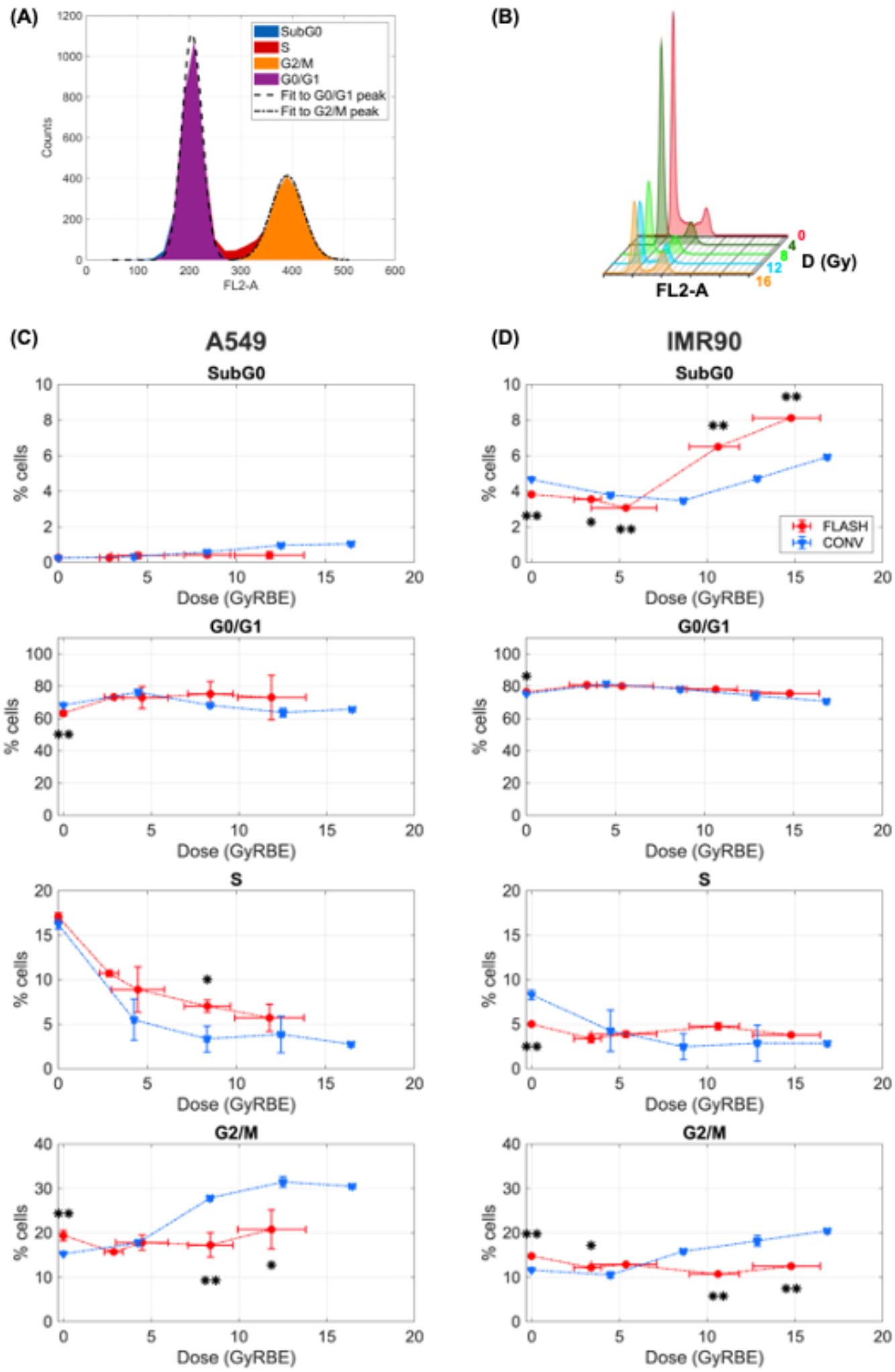
(A) and (B) panels: FL2-A histograms and detail of feature extraction process. (C) and (D) panels: percent of cells in each phase of the cell cycle for FLASH (red) and conventional (blue) irradiation, as determined by flow cytometry, for A549 (C) cells and IMR90 (D) cells. Dashed lines connecting the markers are provided only to guide the eye. Two-tailed P-values were calculated from the Z statistic and are shown with stars in the plots: (*) = p<0.05; (**) = p<0.01.

### 3.4 Expression of p21

Lastly, we also analyzed via immunofluorescence the expression of the cyclin-dependent kinase (CDK) inhibitor p21 which promotes cell cycle arrest in response to a variety of stimuli (Karimian 2016). The experimental data (Figure 6) were fitted to the following expression, representing the fraction of cells positive for p21, *P*_*p*21_(*D*), after receiving a dose *D*.

**Figure 6.**
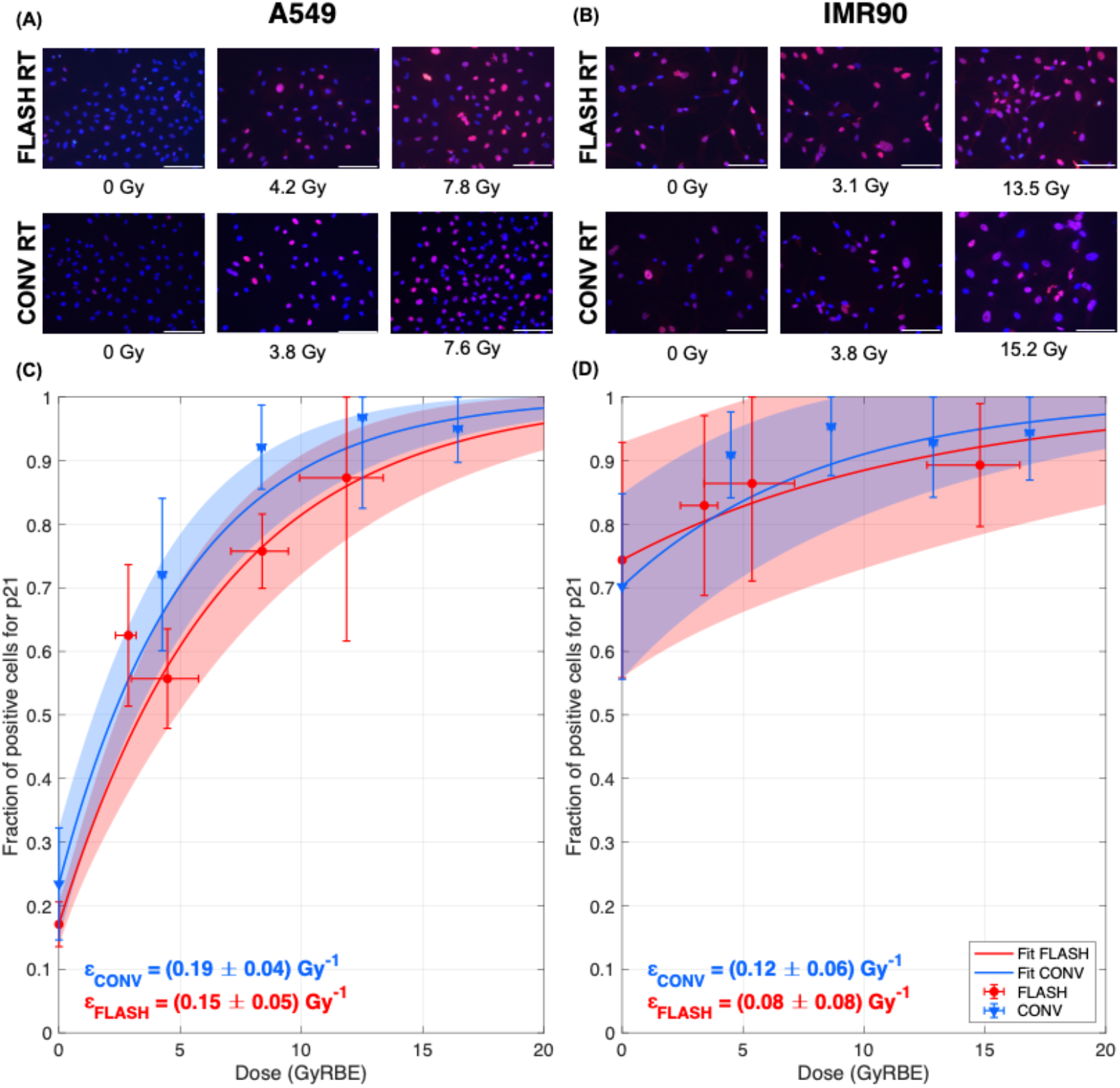
(A) and (B) panels: Fluorescence images of cells A549 (A) and IMR90 (B) expressing p21 (red) merged with the DAPI staining (blue). (C) and (D) panels: fraction of cells labeled positive in IF assay for p21 expression for FLASH (red) and conventional (blue) irradiation, in A549 cells (C) and IMR90 cells (D). Shaded areas indicate 1-sigma confidence intervals for the fit to equation [1]. Scale bar of 100 μm.

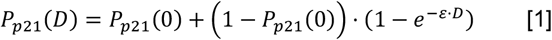

For A549 cells we observed dose-dependent increase in p21 expression. However, no significant differences (at 95% confidence) were observed between conventional and FLASH dose rates, although a tendency to induce higher levels of p21 with the conventional dose rate was found (Figure 6A) which might correlate with the arrest in G2/M observed (Figure 5A), particularly at 8 Gy (p∼0.08). Regarding the healthy fibroblasts, which express higher basal levels of p21, no significant increase in its expression was observed either with conventional or FLASH irradiation (Figure 6B).

## 4. Discussion and conclusions

Our work demonstrated the feasibility of delivering biologically relevant doses at FLASH rates using the synchrocyclotron room of a clinical proton accelerator to small biological samples. Using the appropriate scatterer and positioning system, we administered known doses at over 1000 Gy/s to Eppendorf vials containing pellets of cancer and healthy cell lines. The repeatability of the irradiations in this first experiment, including the positioning uncertainty, falls within a 10% margin, which could be improved in subsequent irradiations. The amount of radiation received by the experimenters is lower than that experienced during a transatlantic flight and is compatible with the regular conduct of experiments, provided that good radiation protection practices are adhered to, and specific radioprotection protocols are implemented for sample positioning and personnel monitoring. Using an automatic positioning and imaging system (i.e. a 2-axis robot) would improve positioning accuracy and allow for irradiation of larger items using a controlled superposition of spots.

Radiochromic films provided an accessible, dose-independent passive dosimeter for this experiment. However, due to the processing time of the films, the exact dose distributions in the samples could not be known immediately. Since commissioning and irradiation experiments were performed in a single shift, it was therefore not possible to replicate the exact same doses for both FLASH and conventional irradiation, making it harder to compare the results of both branches.

Regarding the biological results, we are aware that, in our study, FLASH and conventional irradiations were performed at different parts of the proton irradiation field, and therefore at different relative biological effectiveness (RBE). While we used the available models to calculate RBE-weighted doses and have used those to compare our data, RBE is an endpoint-dependent quantity, derived from survival curves, and cannot certainly capture all possible radiation damage effects. However, these RBE effects are not expected to affect the conclusions drawn from our cell cycle arrest data, as the variation in the fraction of cells in arrest remains consistent across a broad range of doses, substantially exceeding the variability expected from extreme RBE values. Indeed, we performed a sensitivity analysis, increasing the uncertainty in all RBE values to 0.1(see Figure S1), and the conclusions of our study remained unchanged: the survival assays for the IMR90 (Figure 4) did not show a significant difference between the FLASH and conventional arms (the result of the p-value test when comparing the LQ α parameter decreased to p∼0.58 when considering this extra uncertainty); the fit parameters and their uncertainties shown in Figure 6 remain unchanged, and the significance of the observed FLASH vs. CONV differences in cell cycle arrest (depicted in Figure 5) was maintained. While LET and dose-rate effects are, in fact, impossible to decouple in our experiment, we strongly believe that our results indicate an intrinsic biological difference in radiation damage between FLASH and conventional rates that is worth further investigation. Naturally, for further studies at our irradiation station, equivalent conventional irradiations will be performed in transmission at the entrance channel of the peak, rather than in the SOBP.

The obtained biological results add to the yet scarce body of data for radiobiological study of the FLASH effect in *in vitro* models. To our knowledge, our group is the first to report statistically significant differences between FLASH and conventional irradiations in cell cycle arrest patterns. This, however, does not contradict existing negative evidence, noting that the observed differences in our experiment arise at doses above 7 Gy, and previous research used lower doses: other groups reported no significant effects for HNSCC4, MDA-MB-231, or HeLa cells at 6 Gy (Adrian 2021) or A549 cells at 5 Gy (DelDebbio 2024). However, one study indicated a FLASH effect at 3 Gy for HeLa cells, comparing pulsed (10^9^ Gy/s) and continuous (∼30 Gy/s) low-energy protons at 10 hours post-irradiation; yet, this difference diminished at 24 hours post-irradiation. Given this, future studies should consider earlier time points for assessing cell cycle effects and include doses exceeding 7 Gy.

p21 has been described to modulate radiation responses by regulating cell cycle arrest, apoptosis, DNA repair, senescence and autophagy (Kuang 2021). In our study we observed an increase of p21 levels with conventional dose rates at 8Gy dose in A549, that could correlate with the G2/M arrest observed in this cell line at conventional dose rate that was not observed for FLASH dose rates. p21 is a well-known inhibitor of cell cycle and can arrest its progression in G1/S transition by hampering CDK4,6/cyclin-D or in G2/M transition by repressing CDK2/cyclin-E (Bertoli 2013; Pai 2021). Interestingly, in agreement with our data, in a previous study comparing proton beam irradiation and gamma irradiation, it was shown that proton beam irradiation of A549 cells induced p21 upregulation and G2/M cell cycle arrest (Narang 2015). However, no literature data about p21 regulation using FLASH dose rates could be found. Hence, it would be interesting to explore the correlation between p21 expression and cell cycle arrest patterns comparing conventional and FLASH dose rates in the future.

In terms of survival assays, we do not observe statistically significant difference between FLASH and conventional dose rates for either cell line, as other groups have reported in normoxic conditions. We do, however, see a trend indicating a possible improvement in survival in healthy fibroblasts irradiated under FLASH conditions, in alignment to the reported data with a similar cell line (Adrian 2021). However, care must be taken when comparing cell clonogenic and survival curves, even from the same cell line and oxygenation conditions. For instance, for the A549 cell line, D_10_ values range from 4 Gy (Khan 2021), 8 Gy (Tessonier 2021) and 12 Gy (this work). Similarly, our measured value of D_10_ for the IMR90 is around 15 Gy, contrasting with reported values of ∼4 Gy (Buonanno 2020, Adrian 2021 with MRC-5 fibroblasts). These discrepancies are likely due to differing irradiation protocols, seeding methods, and the usage of cells at late passages, which are more senescent and less radiation-sensitive (Adrian 2021). Therefore, for a fair comparison, efforts must be made to ensure that conventional and FLASH protocols were executed in a precisely identical manner.

## Acknowledgements

This work was funded by Comunidad de Madrid, Spain under projects PRONTO-CM (B2017/BMD-3888) and ASAP-CM (S2022/BMD-7434). We acknowledge support by UCM under project pFLASH (PR27/21-014), and by Spanish MCIN/AEI/10.13039/501100011033 under grants RADFLAP (PID2021-124094OA-I00), FLASHonCHIP (PLEC2022-009256) and PROTOTWIN (TED2021-130592B-I00) and the Investigo program (CT19/23), financed in part by European Union NextGenerationEU/PRTR. The authors acknowledge the support of IBA technical staff at Quironsalud Proton Therapy Center. This is a contribution for the Moncloa Campus of International Excellence, “Grupo de Física Nuclear-UCM”, Ref. 910059.

