## Supplementary materials for "*In vitro* FLASH irradiation of A549 lung cancer cells and IMR90 healthy human fibroblasts in the synchrocyclotron room of a clinical proton therapy system"

### Calculation of radiobiological effectiveness (RBE) for each data point

The McNamara model (McNamara, 2015) was used to estimate a value of radiobiological effectiveness for each datapoint in our study and therefore correct for the different position in the irradiation field of the FLASH and conventional arms, as shown in Figure S1. Table S1 details all RBE values for each datapoints in the study along with their uncertainties.

The following formula was used:

$$RBE\left(D_p, \left(\frac{\alpha}{\beta}\right)_x, LET_d\right) = \frac{1}{2D_p}\left(\sqrt{\left(\frac{\alpha}{\beta}\right)_x^2 + 4D_p\left(\frac{\alpha}{\beta}\right)_x\left(p_1 + \frac{p_2 \cdot LET_d}{\left(\frac{\alpha}{\beta}\right)_x}\right) + 4D_p^2\left(p_3 - p_4 LET_d\sqrt{\left(\frac{\alpha}{\beta}\right)_x}\right)^2} - \left(\frac{\alpha}{\beta}\right)_x\right)$$

Where $D_p$ is the proton dose, $\left(\frac{\alpha}{\beta}\right)_x$ is the literature parameter for photon reference radiation for A549 and IMR90 cells, and LETd is the dose-averaged LET in the sample, calculated with Monte Carlo code TOPAS (Perl 2012). Equation parameters take the following values: $p_1 = 0.999064$; $p_2 = 0.35605$; $p_3 = 1.1012$; $p_4 = 0.0038703$. Uncertainties were estimated for each point by propagating dosimetric uncertainties in $D_p$, chosen uncertainties for $\left(\frac{\alpha}{\beta}\right)_x$ and the standard deviation in LETd in the biological samples calculated with TOPAS (Perl, 2012). A bootstrapping Monte Carlo uncertainty propagating algorithm was employed.

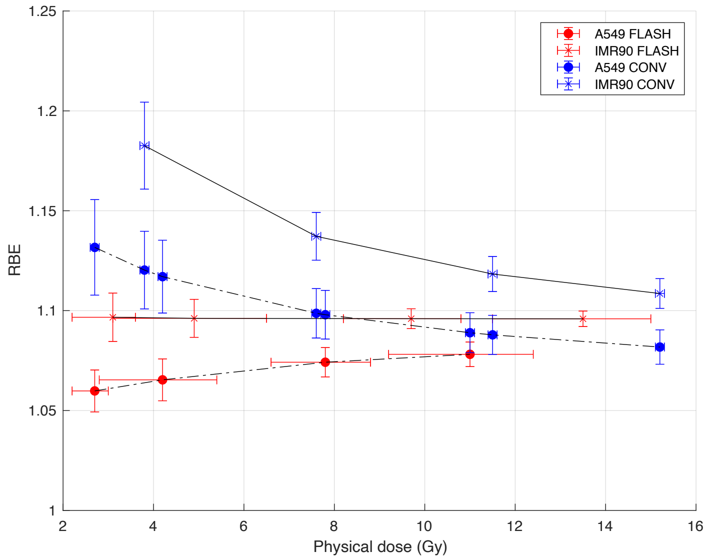


**Figure S1**. RBE values calculated with McNamara model. Error bars indicate standard errors calculated with bootstrapping for each point. Lines connecting the points are provided only to guide the eye.

$LET_{FLASH} = (0.8 \pm 0.1)$ keV/µm

$LET_{CONV} = (4.1 \pm 0.4)$ keV/µm

$\left(\frac{\alpha}{\beta}\right)_{X,A549} = (8 \pm 2)$ Gy

$\left(\frac{\alpha}{\beta}\right)_{X,IMR90} = (3 \pm 1)$ Gy

**Table S1**. Physical and RBE-weighted doses and uncertainties for each point used in the study.

| Series | Physical dose [Gy] | RBE (McNamara model) | RBE-weighted dose [GyRBE] |
|---|---|---|---|
| A549 FLASH | $2.4^{+0.3}_{-0.5}$ | $1.060 \pm 0.011$ | $2.9^{+0.3}_{-0.5}$ |
| | $4.2^{+1.2}_{-1.4}$ | $1.065 \pm 0.011$ | $4.5^{+1.3}_{-1.5}$ |
| | $7.8^{+1.0}_{-1.2}$ | $1.074 \pm 0.007$ | $8.4^{+1.1}_{-1.3}$ |
| | $11.0^{+1.4}_{-1.8}$ | $1.078 \pm 0.006$ | $11.9^{+1.5}_{-1.9}$ |
| IMR90 FLASH | $3.1^{+0.5}_{-0.9}$ | $1.097 \pm 0.012$ | $3.4^{+0.5}_{-1.0}$ |
| | $4.9^{+1.6}_{-1.8}$ | $1.096 \pm 0.010$ | $5.4^{+1.8}_{-2.0}$ |
| | $9.7^{+1.1}_{-1.5}$ | $1.096 \pm 0.005$ | $10.6^{+1.2}_{-1.6}$ |
| | $13.5^{+1.5}_{-2.0}$ | $1.096 \pm 0.004$ | $14.8^{+1.6}_{-2.2}$ |
| A549 CONV | $2.7 \pm 0.1$ | $1.132 \pm 0.024$ | $3.06 \pm 0.13$ |
| | $3.8 \pm 0.1$ | $1.121 \pm 0.020$ | $4.26 \pm 0.13$ |
| | $4.2 \pm 0.1$ | $1.117 \pm 0.018$ | $4.69 \pm 0.14$ |
| | $7.6 \pm 0.1$ | $1.099 \pm 0.012$ | $8.35 \pm 0.14$ |
| | $7.8 \pm 0.1$ | $1.098 \pm 0.012$ | $8.57 \pm 0.15$ |
| | $11.0 \pm 0.1$ | $1.089 \pm 0.010$ | $11.98 \pm 0.16$ |
| | $11.5 \pm 0.1$ | $1.088 \pm 0.010$ | $12.51 \pm 0.16$ |
| | $15.2 \pm 0.1$ | $1.082 \pm 0.009$ | $16.45 \pm 0.17$ |
| IMR90 CONV | $3.8 \pm 0.1$ | $1.183 \pm 0.021$ | $4.49 \pm 0.14$ |
| | $7.6 \pm 0.1$ | $1.137 \pm 0.012$ | $8.64 \pm 0.15$ |
| | $11.5 \pm 0.1$ | $1.118 \pm 0.009$ | $12.86 \pm 0.15$ |
| | $15.2 \pm 0.1$ | $1.109 \pm 0.007$ | $16.85 \pm 0.16$ |
